# RCANE: A Deep Learning Algorithm for Whole-genome Pan-Cancer Somatic Copy Number Aberration Prediction using RNA-seq Data

**DOI:** 10.1101/2024.11.03.621681

**Authors:** Changhao Ge, Xiaowen Hu, Lin Zhang, Hongzhe Li

## Abstract

Transcriptome sequencing (RNA-seq) is widely used in cancer research to study the transcriptome and its role in disease progression. Somatic copy number aberrations (SCNAs) are key drivers of cancer development, and inferring SCNAs from RNA-seq data can provide critical insights for disease classification and treatment prediction. We introduce RCANE, a deep learning-based method designed to predict genome-wide SCNAs across various cancer types using RNA-seq data. RCANE is trained on data from The Cancer Genome Atlas (TCGA) and DepMap cancer cell lines, demonstrating superior performance compared to existing methods. This scalable approach offers a robust solution for improving SCNA prediction in cancer diagnostics and treatment.

---

Somatic copy number aberrations (SCNAs) are a hallmark of cancer, involving large-scale genomic alterations that drive tumorigenesis and cancer progression by affecting gene dosage and altering the expression of oncogenes and tumor suppressor genes. Detecting SCNAs is essential for understanding cancer biology and developing personalized therapies^1–4^. However, SCNA detection traditionally depends on high-cost, high-depth sequencing techniques, such as whole-genome sequencing (WGS) or whole-exome sequencing (WES). In contrast, mRNA sequencing (RNA-seq) offers a more cost-effective and widely used alternative, reflecting cellular activity directly and making it a key component of multi-omic studies. Developing an algorithm that can accurately predict SCNAs from RNA-seq data would significantly reduce costs, making SCNA detection more accessible. However, the nonlinear and complex relationship between gene expression and genomic alterations, along with factors such as DNA-methylation^5–8^and transcriptional adaptation^9^ that influence the RNA transcription, presents a challenge for accurate SCNA inference from RNA-seq data.

Current SCNA detection tools using RNA-seq data generally fall into two categories: modified segmentation-based methods that modify the existing CNA detection methods developed for array comparative genomic hybridization (CGH)^10^ or SNP array data, and machine learning approaches that uses RNA-seq data as predictors. While CNVkit^11^ is capable of estimating SCNAs from RNA-seq data, it was originally designed for DNA sequencing and thus performs poorly on RNA-seq data. Similarly, CopyKAT^12^ was developed for single-cell RNA and doesn’t generalize well to bulk RNA data. Existing machine learning methods, such as CNAPE^13^, improve SCNA prediction from RNA-seq data but often focus on either gene- or chromosome-level predictions, and miss finer patterns. Additionally, these methods typically require large datasets to train the models for SCNAs, limiting their utility in biomedical research where only a few training samples are available. Many other methods like RNAseqCNV^14^, CaSpER^15^ and SuperFreq^16^ also require knowing B-allele frequency, which is unavailable in RNA-seq studies.

These limitations necessitate novel approaches that effectively utilize RNA-seq data for SCNA prediction. The methods proposed in this paper fills this gap by introducing RNA-seq to copy number aberration neural network (RCANE), a deep learning algorithm designed to predict whole-genome SCNAs from cancer RNA-seq data (Extended Data Table 1). Deep learning with an appropriate architecture to capture various dependency among the data is particularly suited to this problem, as it excels at modeling complex, high-dimensional data^17^. In addition, it can fine-tune across diverse datasets, enhancing its generalizability to new studies.

RCANE preprocesses raw mRNA-seq and SCNA intensity data (e.g., the log_2_ of target to reference ratio from the SNP array platform) before neural network modeling (Figure 1a). mRNA-seq data is normalized using transcript per million reads (TPM), with low-expression genes removed. The remaining genes are reordered based on their genomic positions. Since SCNAs typically span large regions and affect hundreds of genes, adjacent genes are grouped into segments. The data is then reshaped into a 3D tensor, where the segments form the last dimension. The log-intensity values of SCNA data are summarized within the same segments. Positive and negative segment graphs are constructed from correlation matrices of segment intensity, where correlations above 0.1 or below -0.1 define positive and negative edges, respectively. These graphs, specific to each cancer type, capture cross-chromosomal correlations by excluding intra-chromosomal edges.

**Figure 1.**
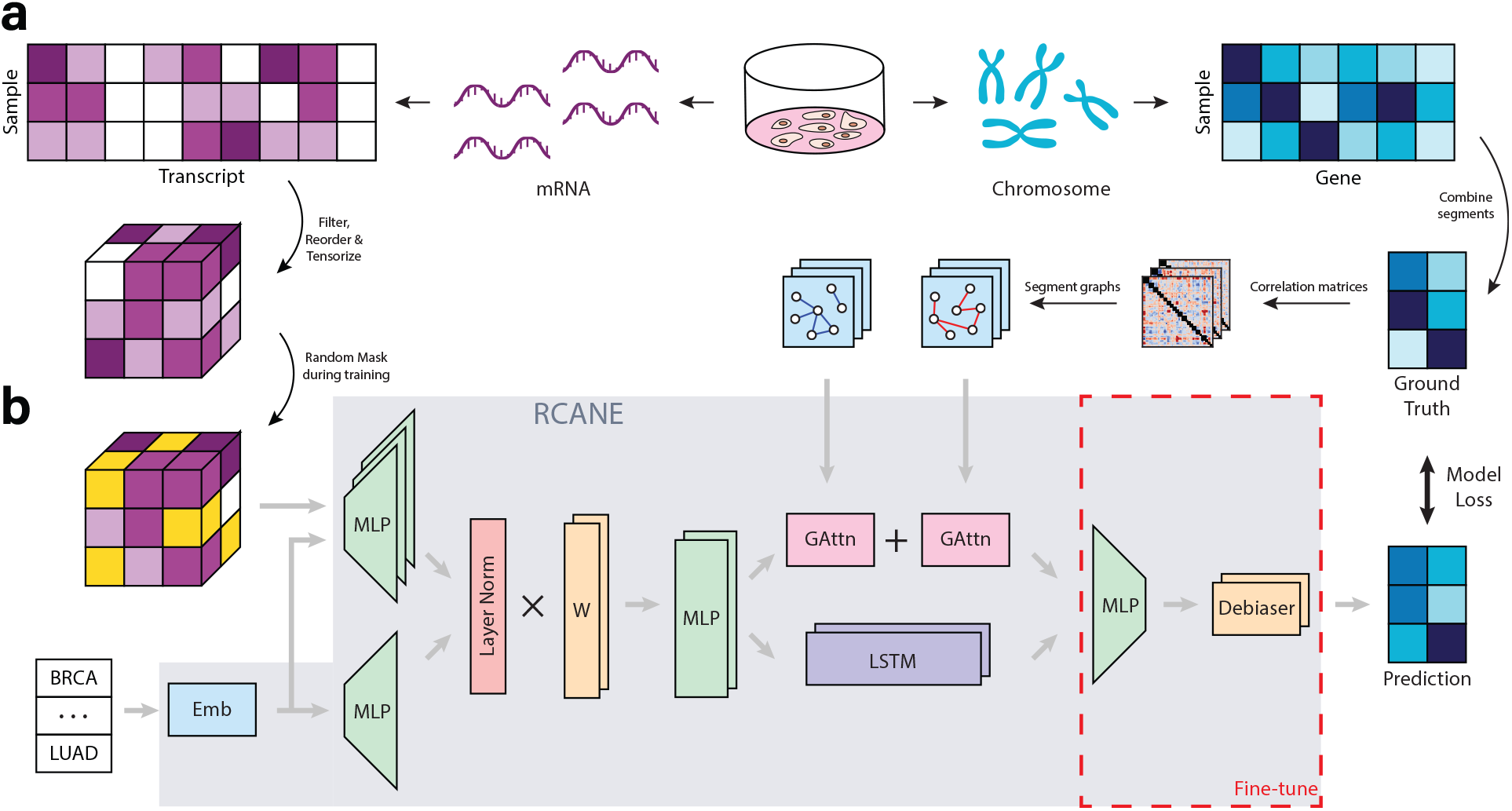
Overview of the RCANE Model. **a**, Data preprocessing. mRNA expression data are filtered and reordered by genomic positions, and gene copy numbers are grouped into segments. These segments are then used to compute correlation matrices and define segment-based graphs. **b**, The RCANE architecture. Cancer types are mapped through embedding. For each segment, mRNA data is adjusted by cancer types and aggregated into a weighted average, which is then processed by an LSTM layer to capturing within-chromosome dependencies and two graph attention mechanisms to capture cross-chromosome correlations. The outputs are merged via an MLP and debiased for each segment. The model is trained using regression loss to align with the ground truth, with fine-tuning applied in the last two layers.

The core architecture of RCANE integrates sequence models and graph neural networks (Figure 1b). To enable the model to learn both the effects of individual gene expression and the relative importance of different genes, a portion of the gene expression values is randomly masked at the start of each epoch. Cancer types are forwarded to an embedding layer, allowing RCANE to capture information across different cancers. Given that various cancers exhibit distinct RNA expression patterns, the unmasked expressions are adjusted using the cancer type embeddings and processed through a multi-layer perceptron (MLP). Outputs from genes within the same segment, combined with cancer type embeddings, are then passed through a layer normalization^18^ step to ensure a mean of 0 and a variance of 1. These normalized values are merged using a weighted average and processed by another MLP. The resulting values are then fed into a chromosome-specific Long Short-Term Memory (LSTM) layer^19^ and two Graph Attention (GAttn) mechanisms^20^, which leverage predefined positive and negative correlation graphs. The LSTM captures both short- and long-range dependencies in gene expression within chromosomes, while the GAttn layers detect cross-chromosomal SCNA patterns, such as the 1p/19q co-deletion in glioma^21^. Finally, the two representations are combined and passed through a debiasing layer for fine-tuning. The model is trained by minimizing the mean squared error between the predicted and actual SCNA intensity values.

We trained and evaluated RCANE using data from The Cancer Genome Atlas (TCGA) data^22^, which includes 33 cancer types of various sample sizes. For external validation, we used an independent dataset of human cancer cell lines representing 17 cancer types from the DepMap^23^ project (Extended Data Figure 1a). Since the cancer cell line data has higher tumor purity resulting in a higher frequency of SCNAs (Extended Data Figure 1b), we applied a correction to the mRNA data to account for these distribution shifts before feeding it into the model (Extended Data Figure 1c, Methods).

To assess the contributions of different model components, we performed an ablation study, focusing on the LSTM, GAttn, and debias layers (Figure 2a). Removing either the LSTM or GAttn resulted in a 7%-9% reduction in MCC on TCGA data and an 11%-14% reduction on cell line data, with accuracy dropping by 2%-3% and 6%-7%, respectively. While the model without the debias layer performed similarly to the full model on TCGA data, it exhibited poorer performance on the cell line data with fine-tuning. This result underscores the importance of including each component in the model architecture.

**Figure 2.**
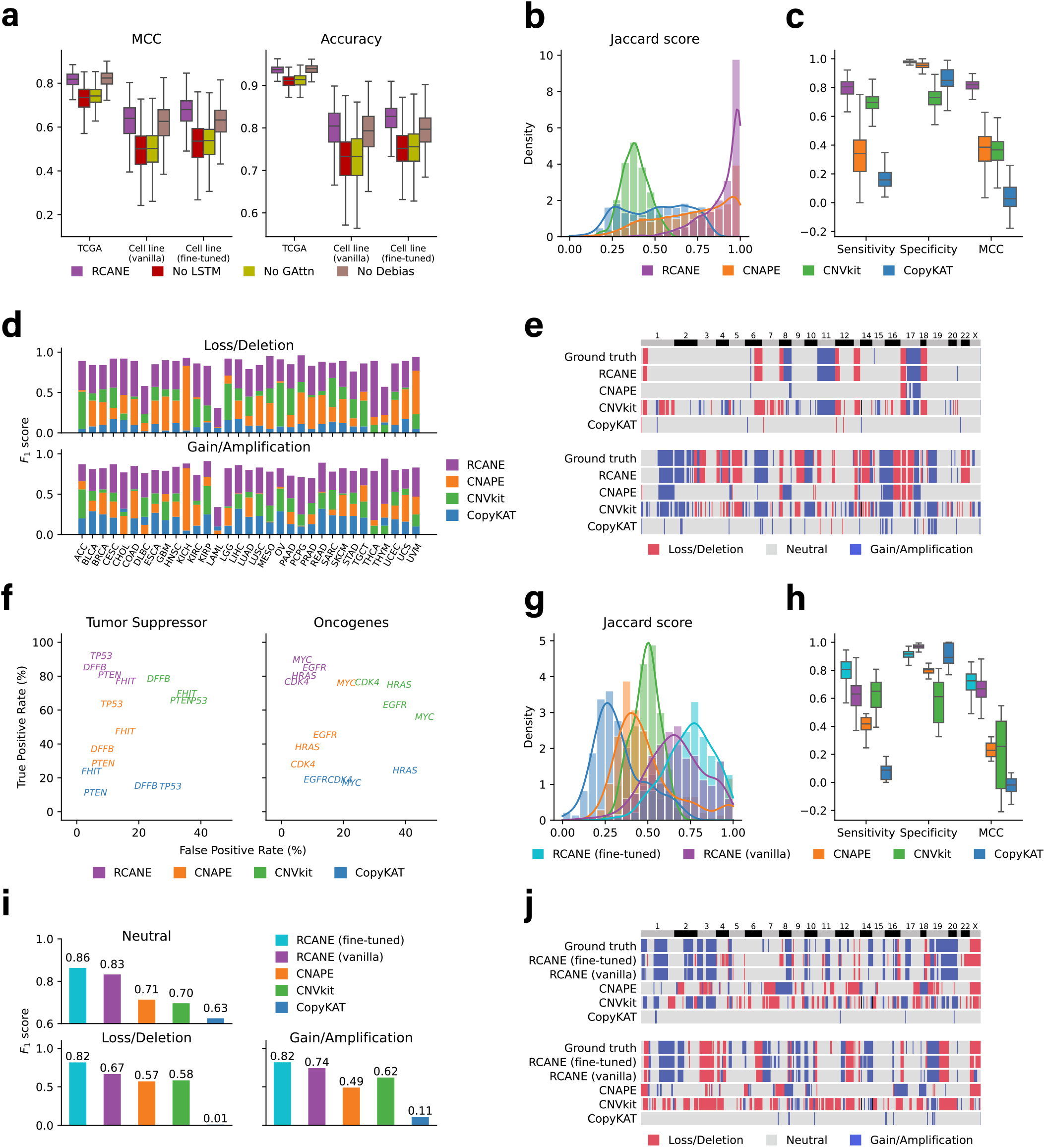
Performance Evaluation of RCANE using TCGA and DepMap cell Line testing samples. **a**, Ablation study comparing MCC (left) and accuracy (right) after removing LSTM, GAttn, and Debias layer, against the full network. **b-f**, RCANE vs. three methods on TCGA data. **b**, Histogram of Jaccard score for whole-genome SCNA prediction. **c**, Boxplot of sensitivity, specificity, and MCC for segment-level SCNA detection. **d**, *F*_1_ scores for copy number loss/deletion (top) and gain/amplification (bottom) across 33 cancer types. KICH and LAML results from CNVkit are excluded due to execution errors. **e**, Whole-genome SCNA prediction of two samples compared with ground truth. **f**, FPR and TPR for 4 tumor suppressor genes (left) and 4 oncogenes (right). **g-j**, Fine-tuned RCANE vs. vanilla RCANE trained on TCGA data and three methods. **g**, Histogram of Jaccard score for whole-genome SCNA prediction. **h**, Boxplot of sensitivity, specificity, and MCC for segment-level SCNA detection. **i**, *F*_1_ scores for SCNA prediction across all samples. **j**, Whole-genome SCNA prediction of two samples compared with ground truth.

In TCGA test samples, RCANE outperformed CNAPE, CNVkit, and CopyKAT in whole-genome copy number prediction, achieving the highest Jaccard score (Figure 2b). It also excelled in segment-wise SCNA detection, with an average sensitivity of 0.80, specificity of 0.97, and MCC of 0.79 (Figure 2c). In comparison, CNAPE tended to under-select, and CNVkit over-selected CNAs, resulting in lower MCCs of 0.37 and 0.35, respectively. As a pan-cancer algorithm, RCANE showed outperform other methods across all cancer types (Figure 2d). While CNAPE performed comparably to RCANE in Kidney Chromophobe (KICH), its performance was unstable across other cancer types. CopyKAT, which is designed for single-cell RNA data, performed the worst in this task. Visualizations of two representative samples demonstrated that RCANE accurately recovered both arm-level and focal SCNAs (Figure 2e). It also achieved a high true positive rate (TPR) and low false positive rate (FPR) for key tumor suppressors like *FHIT* and *DFFB*, as well as oncogenes like *MYC* and *EGFR* (Figure 2g). Interestingly, performance diminished for all methods in hematological malignancies such as Acute Myeloid Leukemia (LAML), likely due to the typically low and unstable RNA content in blood cells, making them less suitable for detecting SCNAs (Figure 2d).

We applied RCANE to the DepMap cell line dataset to demonstrate its generalizability. Without fine-tuning, it outperformed other methods in whole-genome SCNA recovery (Jaccard score, Figure 2g) and segment-wise SCNA prediction in sensitivity, specificity and MCC (Figure 2h). The fine-tuned model further improved its sensitivity and enhanced whole-genome SCNA prediction compared to the vanilla version trained only on TCGA data. Due to small sample sizes, *F*_1_ scores were averaged across samples (Figure 2i). The fine-tuned RCANE achieved *F*_1_ scores of 0.86 for segments with neutral copy number, 0.82 for loss/deletion, and 0.82 for gain/amplification), surpassing the vanilla model’s 0.83, 0.67, and 0.74, with both outperforming other methods. Visualizations showed that while the vanilla RCANE underperformed slightly compared to the fine-tuned version, it still captured most of the key features (Figure 2j).

Originally trained on SNP chip intensity ratio data, RCANE extends its application to predicting log-intensity values, offering greater precision compared to categorical SCNAs, enabling more detailed research. RCANE outperformed CNVkit by providing more accurate estimation of the log-intensity ratios (Extended Data Figure 2a, 3), capturing the intricate relationship between mRNA expression and copy number variations within chromosome segments (Extended Data Figure 2b, 4a), leading to a more accurate genome-wide intensity profile prediction (Extended Data Figure 2c, 4b).

Additionally, RCANE’s use of a masking and weighted-average mechanism enables the identification of SCNA-related genes through model-assigned weights, with layer normalization ensuring that gene importance is solely reflected by the weight values (Extended Data Figure 5). This approach also addresses the issue of missing data by masking rather than imputation and enhances model robustness (Extended Data Figure 6). Generally, genes assigned higher weights are more influenced by SCNAs, while those with lower weights are largely independent of SCNA effects (Extended Data Figure 7a). In our model, we identified SCNA-associated genes such as *POLR2H* and *ABCF3*, as well as genes with lower relevance, such as *LINC02069* and *LINC02054* (Extended Data Figure 7b,c).

In summary, RCANE provides a cost-effective solution for predicting SCNAs from RNA-seq data, serving as a viable alternative to traditional sequencing-based methods. This deep learning model effectively captures the complex relationships between RNA and CNA, outperforming existing tools. Its strong generalization across external datasets enhances its utility in various cancer studies. Additionally, RCANE identifies SCNA-related genes, offering valuable insights into the impact of genomic alterations on gene expression. We anticipate that this model will be widely applicable to other cancer types, with potential future extensions, such as multi-omics integration^24,25^, further broadening its impact in cancer genomics.

## Methods

### RCANE implementation

RCANE was developed using Python (version 3.8.19), NumPy (version 1.26.4), PyTorch (version 2.4.1+cu121), and PyG (version 2.5.3). The input to the neural network model consists of three components: a tensor of dimensions *B* × *N*_*s*_ × *N*_*t*_ representing mRNA expression, a vector of size *B* encoding the cancer type, and a *B*×*N*_*s*_ ×*N*_*t*_ dimensional mask, where *B* is the batch size, *N*_*s*_ = 1514 is the number of segments, and *N*_*t*_ = 20 is the number of transcripts per segment. The masking tensor has two main functions. First, it handles end-of-chromosome masking. If the final segment of a chromosome contains fewer than *N*_*t*_ transcripts, the extra positions in that segment are masked, ensuring that these values are not passed to the neural network. Second, random masking is applied during training to prevent over-fitting and encourage the model to learn the independent effects of SCNA on each transcript. This approach also enables the model to identify the relative importance of each gene: genes with higher learned weights tend to be more strongly associated with SCNA, while those with lower weights are less relevant (Extended Data Figure 7).

Cancer type information is processed through an embedding layer. Since tumor samples from different cancer types are typically collected from different tissues, they may have distinct RNA expression characteristics. These inherent tissue-specific differences are not helpful for SCNA prediction. Therefore, each transcript’s value is adjusted based on the cancer type to account for these variations. The adjusted value is then passed through a dedicated MLP, with an input size of 1 and an output size of *N*_out_, resulting in a total of *N*_*s*_ ×*N*_*t*_ MLPs (where *N*_*s*_ is the number of segments and *N*_*t*_ is the number of transcripts per segment). Given that different cancer types have distinct SCNA baselines, their embeddings are also processed through an MLP with the same output size as the input for the next layer. For each segment, the *N*_*s*_ × *N*_*t*_ output vectors from the transcripts, along with the cancer type output vector, are combined into a tensor of size *N*_*s*_ × (*N*_*t*_ + 1) ×*N*_out_.

As the magnitude of different transcripts can vary greatly during training, we apply layer normalization along the last dimension to normalize their scales. To combine the information from each transcript, a weighted average of the rows in this matrix is computed using a weight vector of size *N*_*t*_ + 1, transformed by the softmax function:

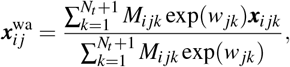

where 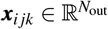 represents the output vector of the *i*th sample, *j*th segment and *k*th transcript (or the cancer type when *k* = *N*_*t*_ + 1), *M*_*i jk*_ ∈ {0, 1} is the masking value with *M*_*i jk*_ = 0 indicating the value is masked, and 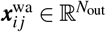 is the resulting weighted average output. The weight *w*_*jk*_ ∈ ℝ reflects the importance of each transcript in predicting the copy number of the *j*th segment. The resulting output is then passed through segment-specific MLPs, followed by the Graph Attention and LSTM layers.

The graph attention layer consists of two attention blocks: one models positively correlated segments and the other models negatively correlated segments. Both blocks receive the output from the previous MLP layer as node features but operate over different sets of graph edges. The first block processes edges defined by positive correlations, while the second processes edges defined by negative correlations. These two correlation types are modeled separately, as their inherent differences require distinct sets of model parameters.

In parallel to the graph attention layers, the model also incorporates an LSTM layer composed of 23 LSTM units, each corresponding to a chromosome. Within each LSTM sequence, individual cells represent chromosome segments. Each cell receives input from the segment-specific MLP and outputs neighborhood-corrected information for the corresponding segment. This design enables the LSTM layer to capture sequential dependencies across segments within each chromosome. To integrate sequential and relational information, the outputs from both the LSTM layer and the two graph attention blocks are concatenated. This combined representation is then passed through a universal MLP shared across all *N*_*s*_ segments, which generates a scalar output for each segment.

The final component of RCANE is a debiasing block, designed to correct biases during fine-tuning. When applied to new data, such as cancer cell lines, variations in characteristics like tumor purity can impact the model performance. To mitigate this, the debiasing block is constructed using *N*_*s*_ fully univariate neural networks, where each layer, including the hidden layers, is univariate. The output of this block represents the predicted copy number. The model is trained by minimizing the mean squared error between the predicted and true copy number intensities across all segments. Other regression loss criteria such as Huber loss are also implemented for future analysis.

### Missing RNA-seq data

In many cases, RNA data of interest may contain missing values compared to the data used to train the RCANE model, often due to differences in data sources or varying platforms for data generation^26^. Missing data presents a significant challenge in machine learning, particularly in deep learning, as it can severely impact model accuracy and robustness.

However, RNA data is known to be highly correlated and often resides on a low-dimensional manifold^27,28^. For instance, genes within the same biological pathway often exhibit co-expression patterns^29,30^, meaning that missing values for some genes may not result in substantial information loss. Leveraging this property of RNA data, RCANE employs a masking and weighted-average mechanism to address missing data issues. During prediction, missing values are masked, and the available RNA data from existing genes within each segment are used to predict SCNAs.

To evaluate RCANE’s performance under missing data conditions, we simulated missing data by randomly deleting a portion of RNA-seq data in both TCGA and cell line datasets. For cell line data, we assessed both the vanilla RCANE model and its fine-tuned version. Within the RNA expression matrix, each element had a defined probability of being missing, with missing probabilities *p* ∈ {0, 0.1, 0.2, 0.3, 0.4, 0.5, 0.6}, where *p* = 0 indicates no missing data. We compared RCANE’s performance with two data imputation methods: KNN imputation with *k* = 5 and median imputation (Extended Data Figure 6). The TCGA data from model training was used for the imputation process. We demonstrated that RCANE’s approach maintained robust performance even when a substantial portion of RNA data was missing, while traditional imputation methods showed significant performance deterioration under the same conditions.

### Steps for data processing

#### TCGA dataset

The TCGA project analyzed over 11,000 tumor samples across 33 cancer types, encompassing both common and rare forms of cancer, such as glioblastoma, breast cancer, lung cancer, as well as rarer types like adrenocortical carcinoma and mesothelioma. Sample sizes varied by cancer type, with hundreds of samples typically analyzed for each. The largest groups included Breast Invasive Carcinoma (BRCA) with over 1,000 samples, Lung Adenocarcinoma (LUAD) and Glioblastoma Multiforme (GBM) with more than 500 samples each, and Colon Adenocarcinoma (COAD) with over 400 samples.

For our study, TCGA data was downloaded using the TCGAbiolinks^31^ (version 2.25.3) through Bioconductor^32^ (version 3.18), which provided both raw and TPM-normalized mRNA expression matrices, along with SCNA data. The mRNA expression data consisted of 60,660 transcripts. Raw mRNA expression data was used as input for CNVkit as recommended^11^, while TPM-normalized data was used for all other methods. For copy number variation analysis, we retrieved Affymetrix Genome-Wide Human SNP Array 6.0 data, which had been transformed to log_2_ ratio and processed using the circular binary segmentation algorithm^33^. Copy number intensities greater than 0.25 were classified as gains or amplifications, while those below -0.2 were classified as losses or deletions. A total of 8,900 tumor samples were used for training and 2,226 for testing, with the data randomly split across cancer types. For methods other than RCANE, separate models were trained for each cancer type to account for the heterogeneity among cancers.

To prepare the input for RCANE, transcripts located on the Y chromosome or expressed in fewer than 30% of the samples were excluded, resulting in a set of 30,096 transcripts. These transcripts were reordered by genomic location across 23 chromosomes based on the genome reference GRCh38. Within each chromosome, 20 adjacent transcripts were grouped into segments, producing a tensor of size *N* × 1514 × 20. The mRNA expression data was then transformed using log_2_(1 + TPM). For copy number data, we generated the RCANE output by calling the copy number for each selected transcript, with each segment represented by the median value of the 20 transcripts within that segment.

To build the correlation graphs used by RCANE, we computed a segment correlation matrix for each cancer type based on copy number intensity data from samples of that cancer type. Positive correlation graphs included edges between segment pairs with correlation values greater than 0.1, while negative correlation graphs included pairs with correlation values below -0.1. Only segment pairs from different chromosomes were considered when constructing the correlation graphs.

#### Cell line dataset

The cell line data was downloaded from the DepMap portal (https://depmap.org/portal/) and the Score project (https://depmap.sanger.ac.uk/). We selected samples with cancer types overlapping those in the TCGA dataset and randomly split them by cancer type, resulting in 114 training samples and 266 testing samples across 17 cancer types. For RCANE, the training samples along with cancer type information were used for fine-tuning. For other methods, the training samples from different cancer types were combined and used for training from scratch due to the small sample size.

The mRNA expression data initially contained 53,961 transcripts, from which we selected 29,088 transcripts that overlapped with the TCGA data. Other missing transcripts were imputed using the 5 nearest neighbors with scikit-learn (version 1.3.2). As the cell line data does not provide raw counts, we used log_2_(1 + TPM) as input for all four methods examined in this work. Due to the significant distribution shift between the TCGA and cell line mRNA expression data, we applied the ComBat^34^ function from sva^35^ (version 3.50.0) to correct for batch effects, using the TCGA data as the reference and cancer type as covariate (Extended Data Figure 1c).

The copy number intensity data for the cell line was processed by the Broad Institute. Unlike the TCGA data, these values were not log_2_ transformed, so we applied the transformation manually. We then summarized the intensities into segments and defined losses/deletions and gains/amplifications in a consistent way as the TCGA data.

### Data analysis

The performance of RCANE and other methods was evaluated using scikit-learn (version 1.3.2). Visualizations are based on pandas (version 2.0.2), matplotlib (version 3.9.1) and seaborn (version 0.13.2). In the box plots, the center line indicates the median, the box limits represent the first and third quartiles, and the whiskers extend to 1.5 times the interquartile range. The Jaccard score was calculated as follows:

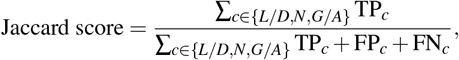

where *L/D, N*, and *G/A* represent copy number loss/deletion, neutral, and gain/amplification categories, respectively. TP_*c*_, FP_*c*_ and FN_*c*_ denote the true positives, false positives, and false negatives for category *c*. The *F*_1_ score for each category is defined as:

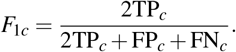

For SCNA detection, we define:

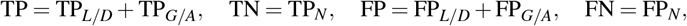

And

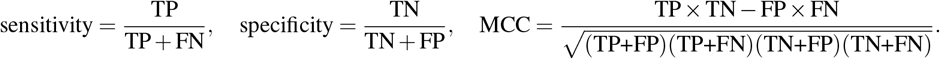

The Pearson’s correlation coefficients (*r*) and the corresponding *p* values were computed in SciPy (version 1.13.1).

## Data Availability

The TCGA and DepMap cell line data used for model training and evaluation were obtained as described in Methods. The preprocessed data, along with model checkpoint files trained on TCGA data and fine-tuned with cell line data, are available in a zipped file at https://doi.org/10.5281/zenodo.13953634.

## Code Availability

The scripts for model training, notebooks for performance evaluation, data analysis, and figure generation are freely available at https://github.com/HowardGech/RCANE, under the open-source MIT license.

---

**Extended Data Figure 1.**
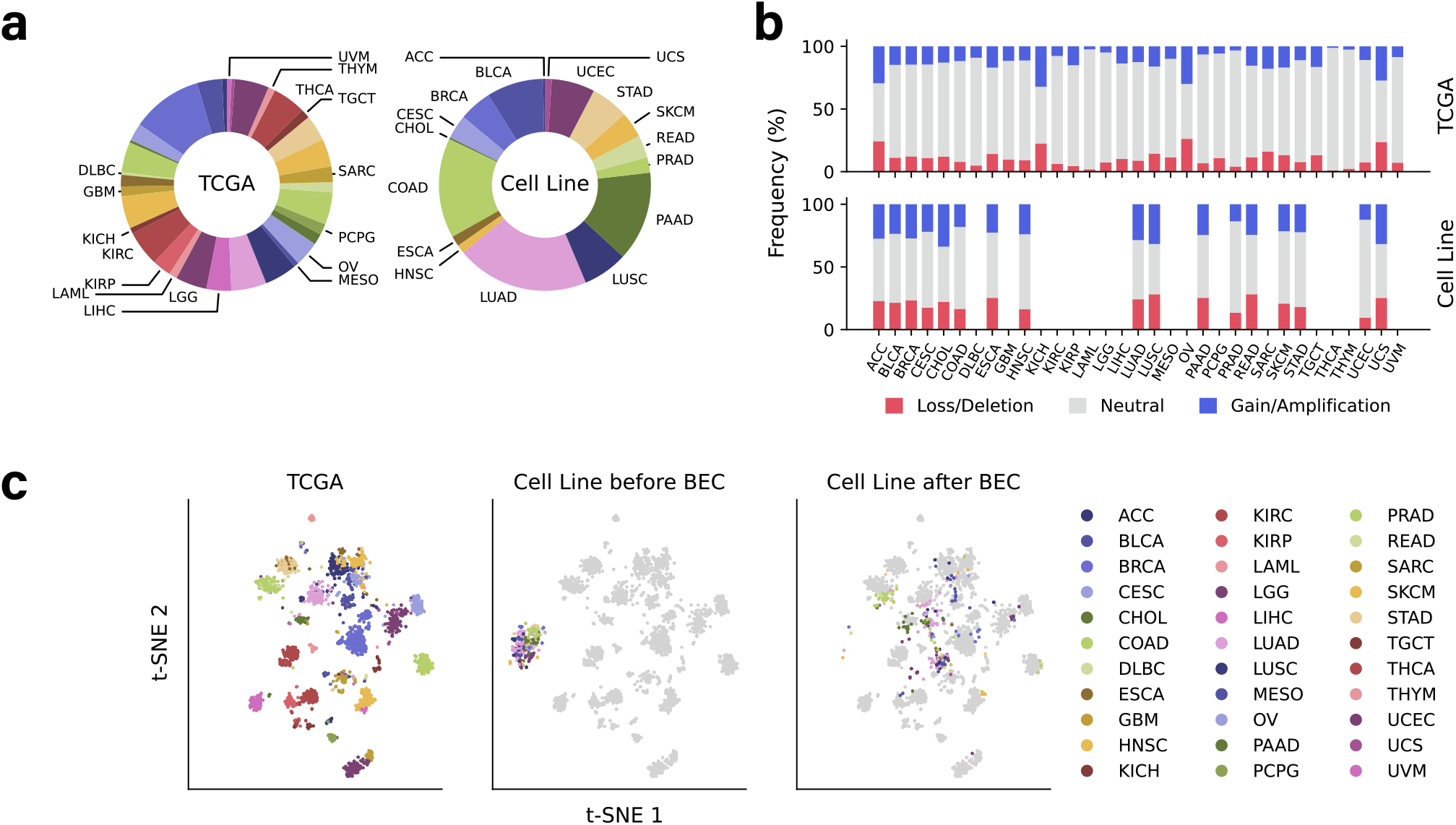
Overview of TCGA and cell line datasets. **a**, Distribution of cancer types in the TCGA dataset (left) and the cell line dataset (right). The TCGA dataset includes 33 cancer types, while the cell line dataset covers 17 of these types. **b**, Frequency of copy number alterations (Loss/Deletion, Neutral, and Gain/Amplification) in the TCGA dataset (top) and the cell line dataset (bottom). **c**, t-SNE visualization of mRNA expression. Left: TCGA data. Middle: cell line data before batch effect correction (BEC). Right: cell line data after BEC.

**Extended Data Figure 2.**
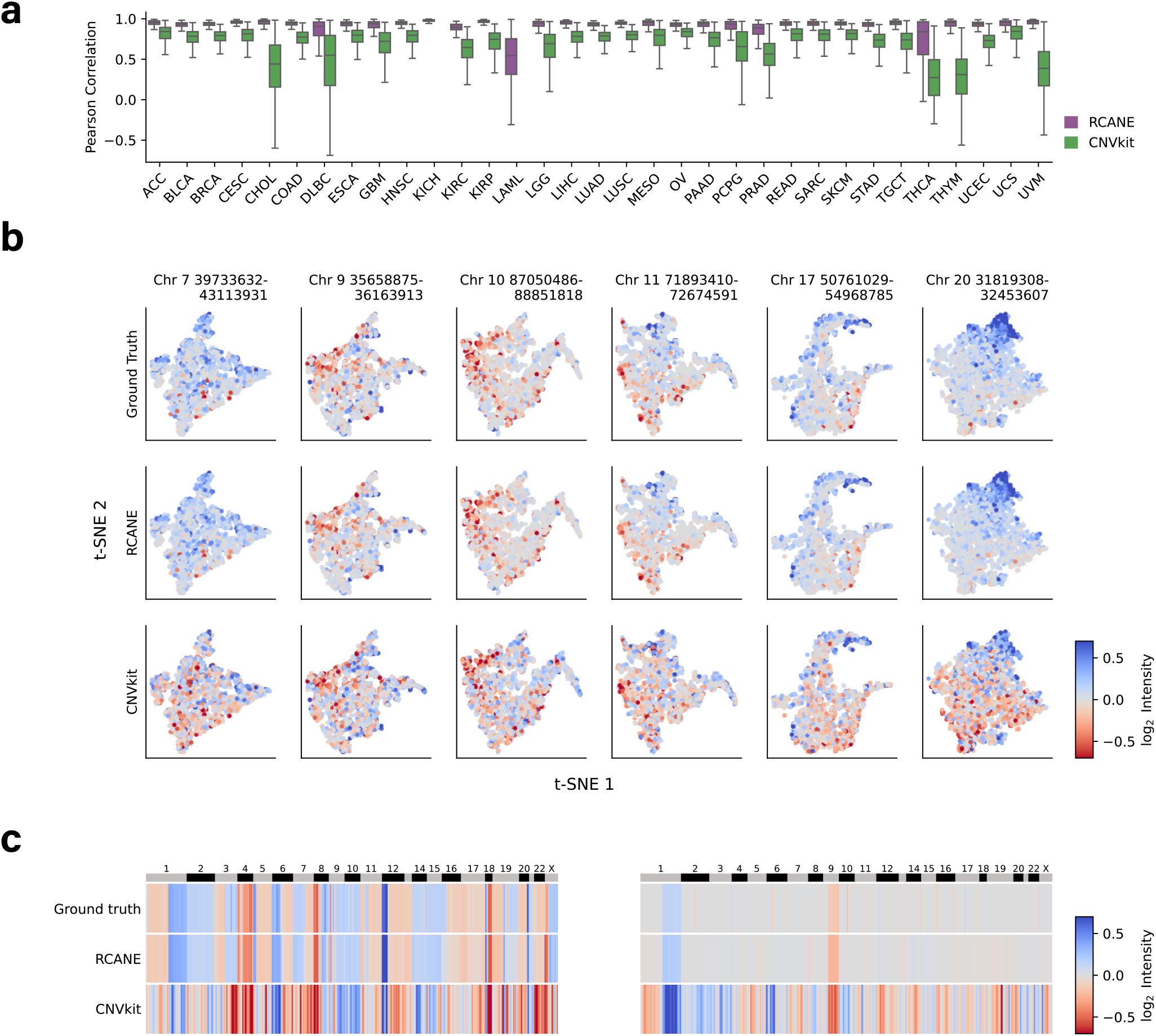
Intensity estimation of RCANE and CNVkit on TCGA testing data. **a**: Boxplot of Pearson correlation between intensity prediction and ground truth for all cancer types. KICH and LAML results from CNVkit are excluded due to errors. **b**, t-SNE plots of ground truth and predicted intensity with dot locations based on mRNA expression data. Colors indicate copy number intensity in log_2_ ratio. **c**, Examples of whole-genome intensity prediction for two samples. Colors indicate copy number intensity in log_2_ ratio.

**Extended Data Figure 3.**
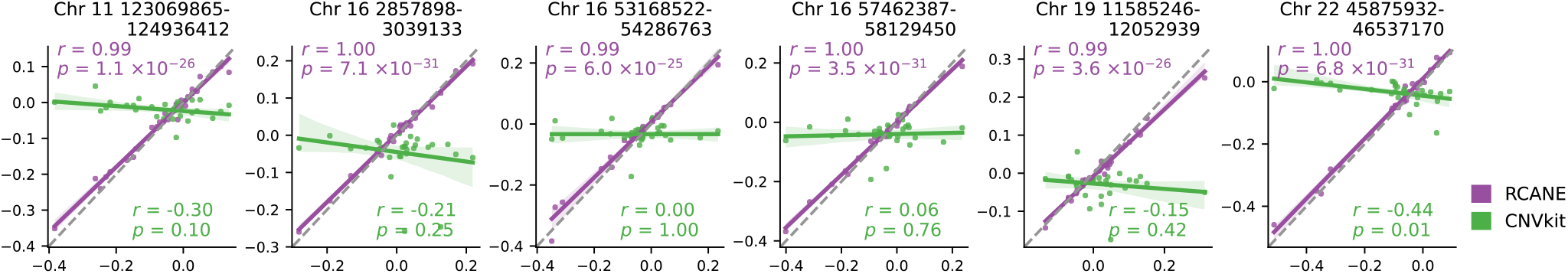
Intensity estimation of selected segments for each cancer types in TCGA samples. Each dot represents one cancer type. Shaded areas represent 95% confidence regions. The *p* value is based on the Pearson’s correlation test.

**Extended Data Figure 4.**
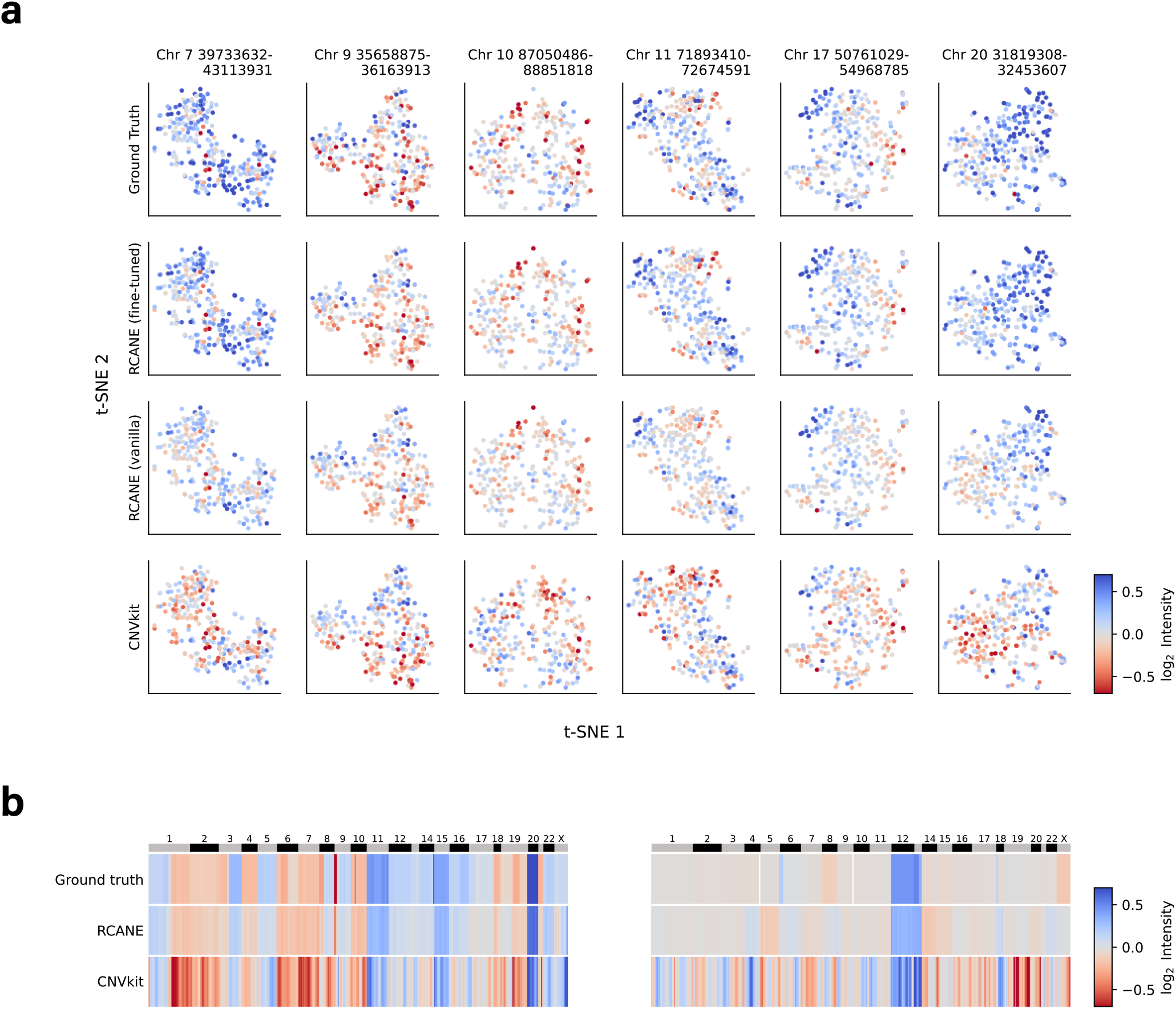
Intensity estimation of RCANE and CNVkit on DepMap cell line testing data. **a**, t-SNE plots of ground truth and predicted intensity with dot locations based on mRNA expression data. Colors indicate copy number intensity in log_2_ ratio. **b**, Examples of whole-genome intensity prediction for two samples. Colors indicate copy number intensity in log_2_ ratio.

**Extended Data Figure 5.**
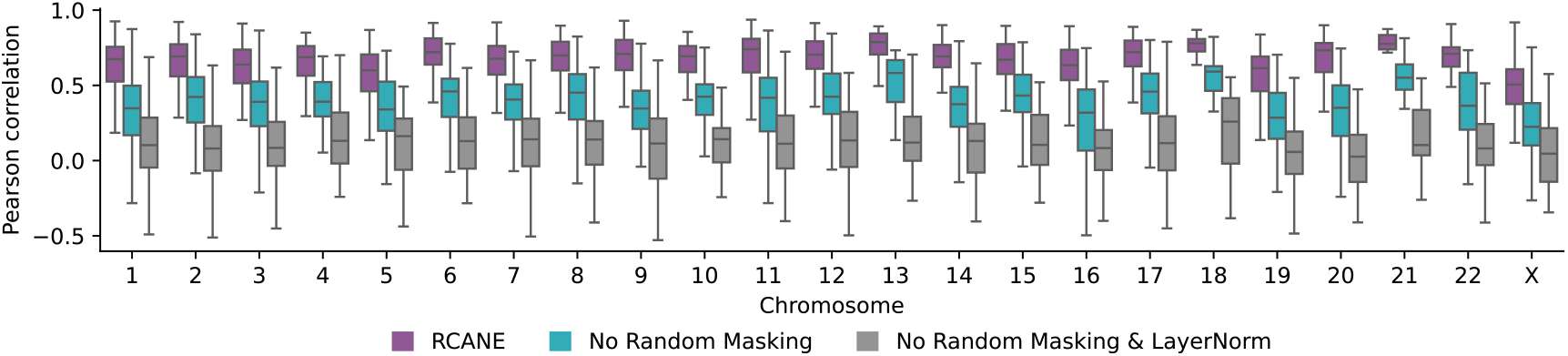
Pearson correlation of gene weight and *r* value between RNA and SCNA data in TCGA samples for 23 chromosomes. Each box is based on the Pearson correlation results of all segments in one chromosome. RCANE: the full RCANE architecture with random masking during model training. No Random Masking: the full RCANE architecture without random masking during model training. No Random Masking & LayerNorm: RCANE model without Layer Normalization mechanism and random masking during model training.

**Extended Data Figure 6.**
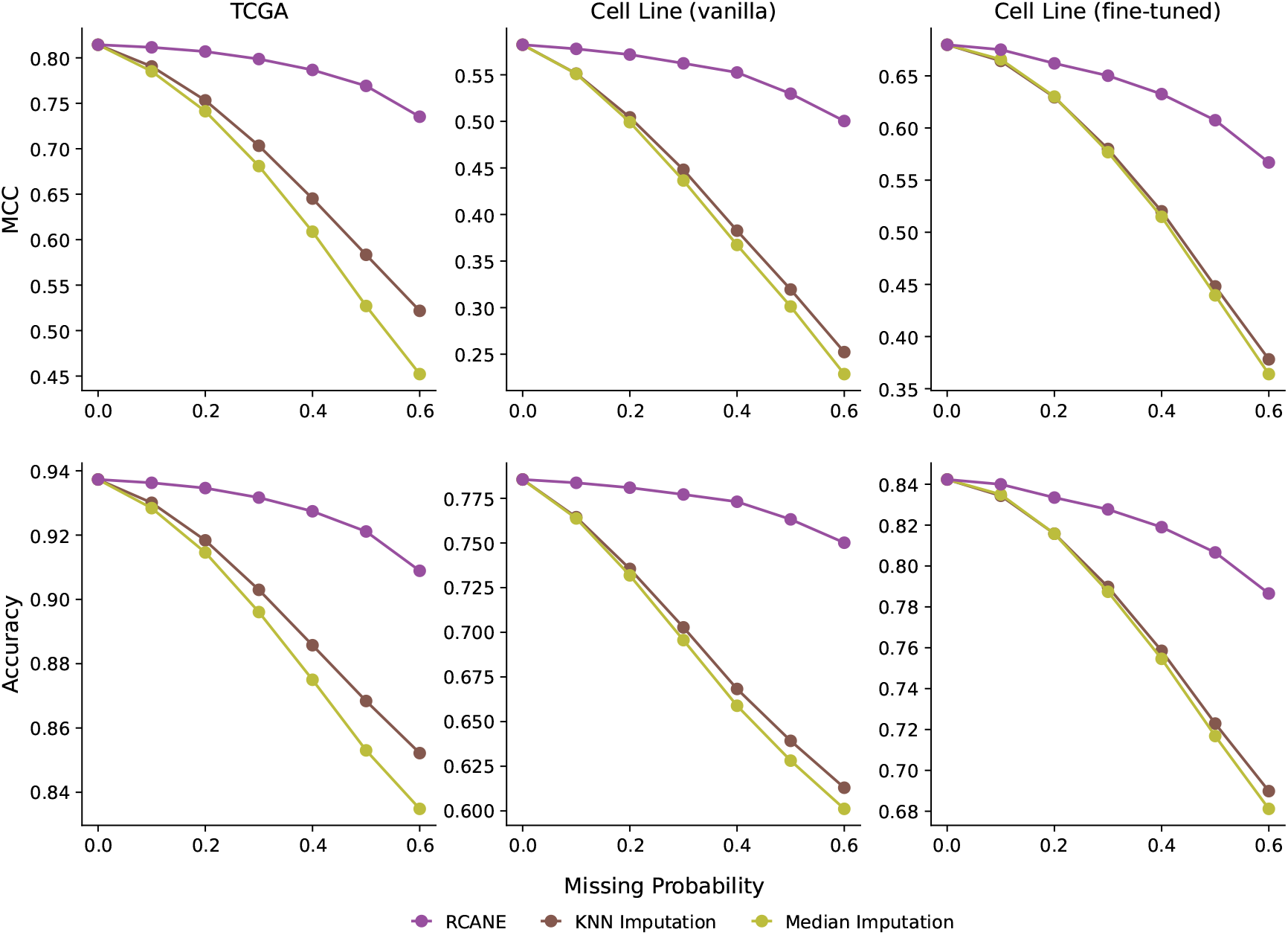
MCC and accuracy of RCANE for SCNA prediction with missing RNA data. RCANE: the RCANE model with missing values masked during prediction. KNN Imputation: the RCANE model with missing values imputed by *k*-nearest neighbors of the same cancer type where *k* = 5. Median Imputation: the RCANE model with missing values imputed by the median value of the same cancer type.

**Extended Data Figure 7.**
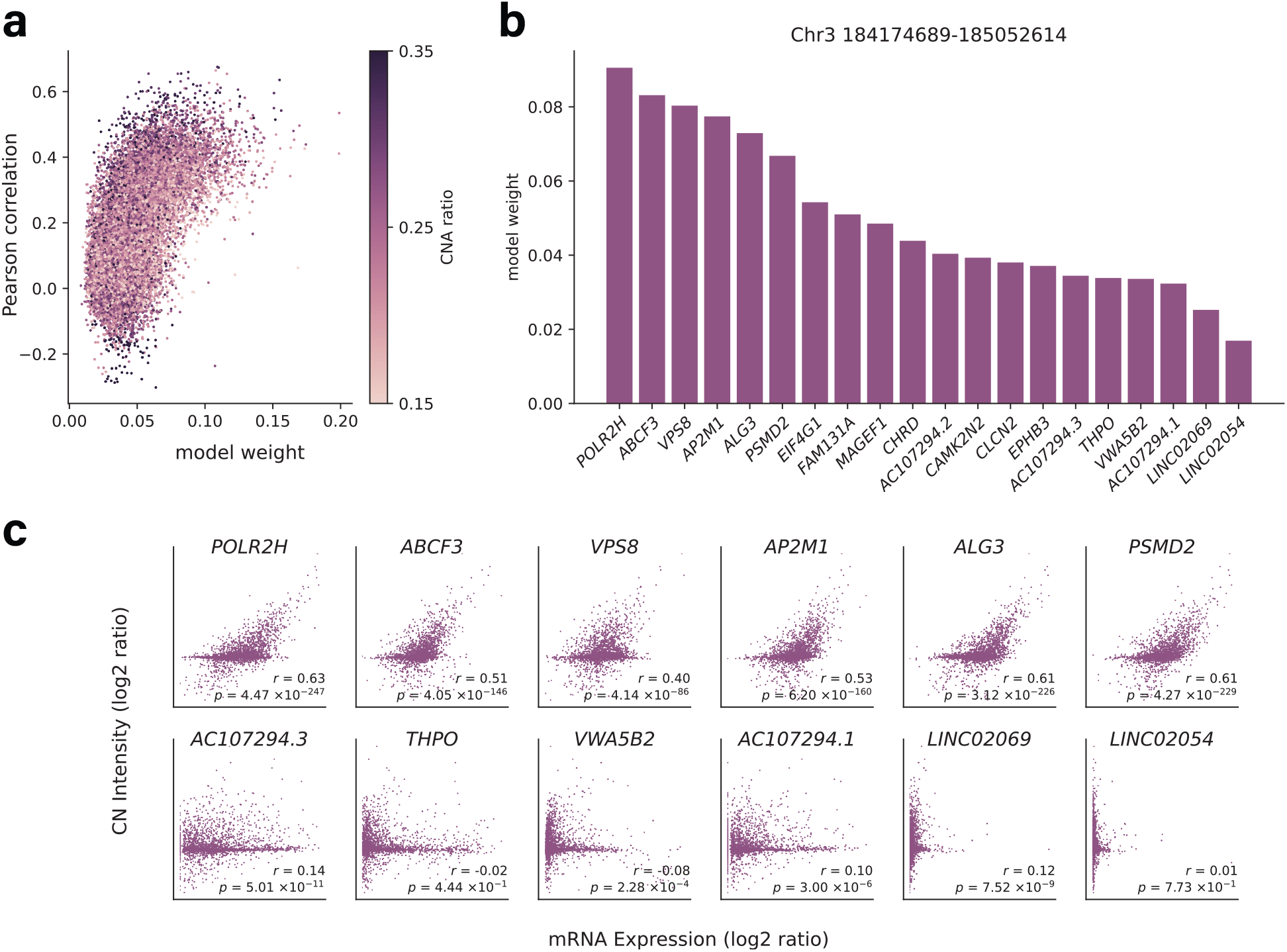
Model weights in RCANE capture CNA-RNA correlations. **a**, Scatter plot showing the relationship between model weights and Pearson correlations of copy number intensity and mRNA expression for each gene. Color indicates the CNA ratio. **b**,**c**, Results of model weights, mRNA expression, and CN intensity for a selected segment. **b**, Model weights of cancer type and 20 genes within one segment. **c**, Scatter plot illustrating mRNA expression and CNA intensity for the genes with the highest (top) and lowest (bottom) model weights, with Pearson’s correlation coefficient (*r*) and its *p* value.

**Extended Data Table 1.** Comparison of RCANE with existing methods for predicting CNA using RNA data.

| Method | Category | Output | Scale | BAF free<sup>a</sup> | Reference free<sup>b</sup> |
| --- | --- | --- | --- | --- | --- |
| RCANE | Deep Learning | Intensity | Whole-genome | ✓ | ✓ |
| CNAPE | Machine Learning | State only | Gene/Chromosome | ✓ | ✓ |
| CNVkit | Segmentation | Intensity | Whole-genome | ✓ | ✗ |
| CopyKAT | Segmentation | Intensity | Whole-genome | ✓ | ✗ |
| RNAseqCNV | Machine Learning | State only | Chromosome | ✗ | ✓ |
| CaSpER | Segmentation | Intensity | Whole-genome | ✗ | ✗ |
| SuperFreq | Segmentation | Intensity | Whole-genome | ✗ | ✓ |

<sup>a</sup> Models that only take RNA expression data as input and don't require B-allele frequencies.
<sup>b</sup> Models that can process the data of a single sample without reference from other samples.

