## Supplementary Material for "RCANE: A Deep Learning Algorithm for Whole-genome Pan-Cancer Somatic Copy Number Aberration Prediction using RNA-seq Data"

### Software workflow and configuration

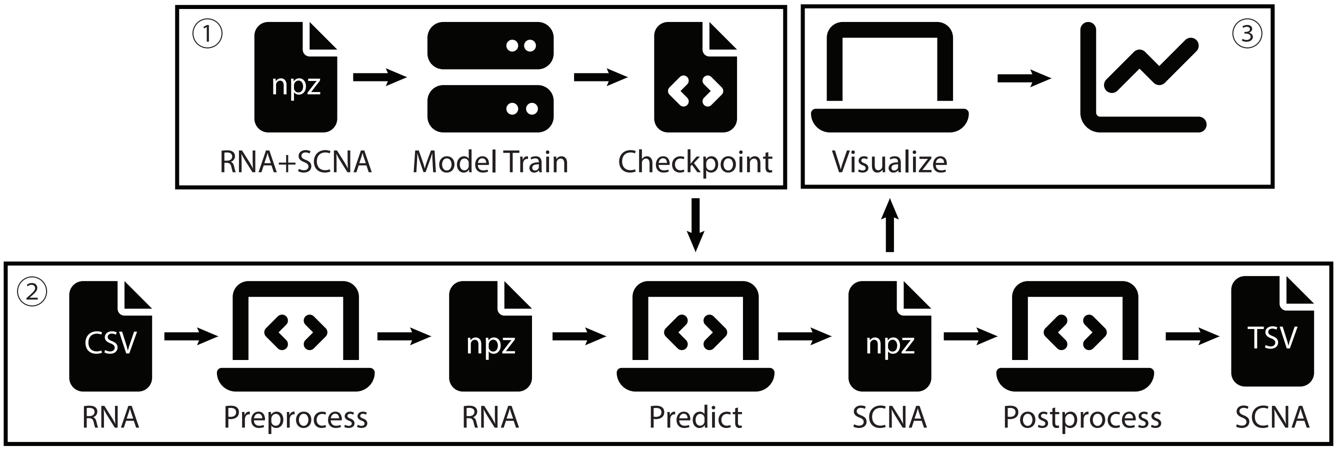


**Figure S.1. The complete RCANE workflow for model training, prediction, and visualization.** The model is trained using both RNA and SCNA data. For prediction, users preprocess their RNA data in CSV format, generating an .npz file that serves as input to the model. The model outputs an .npz file containing predicted SCNAs, which can either be visualized directly or postprocessed into a standard TSV format.

---

The RCANE repository, available at https://github.com/HowardGech/RCANE and https://doi.org/10.5281/zenodo.13953634, enables users to train a new model from scratch, fine-tune an existing model, predict SCNAs from new RNA-seq data using a pretrained model, and visualize predictions. For prediction, the model requires a NumPy zip file (.npz) containing two key elements: a 3D array labeled "rna" for mRNA expression data and a 1D array labeled "cohort" for cancer type information, using TCGA study abbreviations (e.g., "ACC" for Adrenocortical Carcinoma and "BLCA" for Bladder Urothelial Carcinoma). In the RNA data, the first dimension represents samples, the second dimension represents chromosome segments, and the third dimension represents transcripts within each segment. We also recommend including an additional 1D array labeled "ID" to denote sample IDs. The model outputs an .npz file containing the predicted copy number intensity labeled "cna", along with the original "rna" and "cohort".

To create a user-friendly workflow, we provide a preprocessing step that accepts an RNA-seq file in comma-separated values (CSV) format, containing sample IDs, cancer types, and gene names. Additionally, we offer a postprocessing step that generates a tab-separated values (TSV) file for SCNA predictions. This TSV file is similar to a circular binary segmentation (CBS) file and includes the sample ID, chromosome name, segment start and end locations within the chromosome, segment mean value in $\log_2$ scale, estimated segment status (loss/deletion, neutral, or gain/amplification), and the gene names within each segment. The complete pipeline is shown in Figure S.1.

Once the prediction is completed, we provide an interactive tool for visualizing whole-genome SCNA intensities using Plotly (Figure S.2). This tool allows users to select specific samples or chromosomes of interest, or view a combined result for a comprehensive analysis.

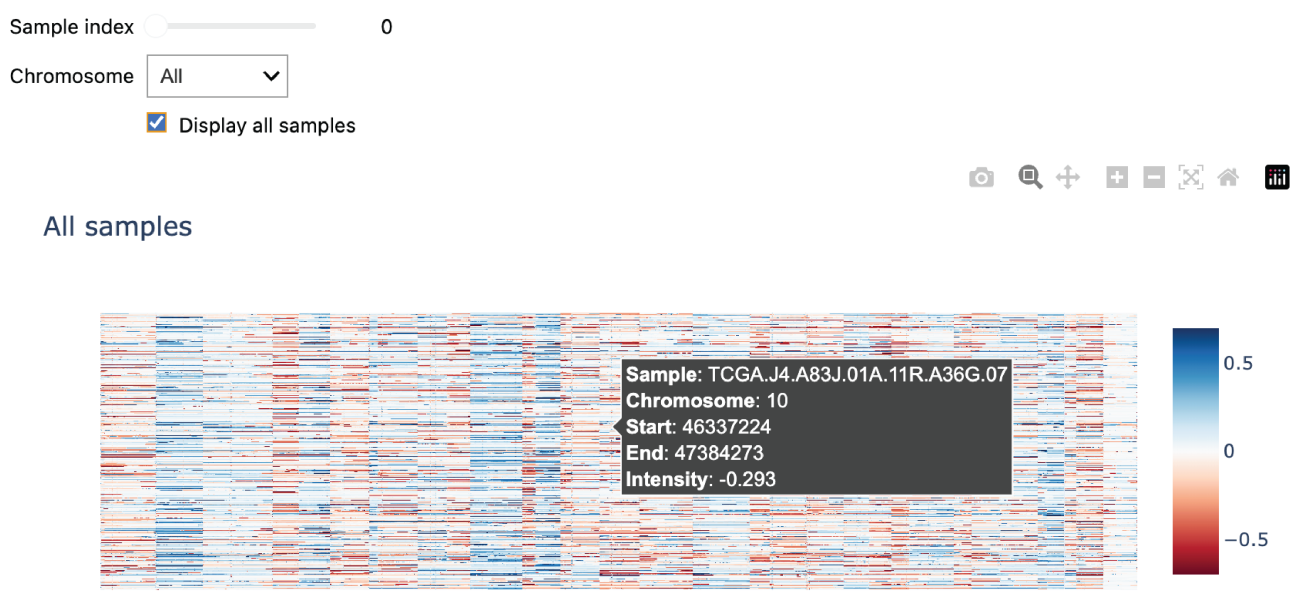


**Figure S.2. Visualization tool for whole-genome SCNAs based on Plotly.** The tool shows the sample ID, chromosome name, start location, end location and intensity value of the SCNAs predicted by RCANE.

Additionally, we provide an interface that allows users to train their own RCANE model or fine-tune a pretrained model. For this, the input .npz file must include an additional 2D array with the same shape as the first two dimensions of the mRNA data, along with the required rna and cohort arrays. For users training a new model from scratch, model parameters—such as the number of layers for different mechanisms, activation functions, and dropout probabilities—can be configured by editing the model parameter YAML file. Since different model architectures may require tailored optimization techniques, users can also modify the training configuration YAML file to adjust settings like the optimizer, learning rate, and number of epochs. In summary, RCANE provides a comprehensive and flexible tool to handle various scenarios, from prediction to model training and customization.
